# Rapid production and recovery of cell spheroids by automated droplet microfluidics

**DOI:** 10.1101/552687

**Authors:** Krzysztof Langer, Haakan Joensson

## Abstract

Droplet microfluidics enables high throughput cell processing, analysis and screening by miniaturizing the reaction vessels to nano- or pico-liter water-in oil droplets, but like many other microfluidic formats, droplet microfluidics have not been interfaced with or automated by laboratory robotics. Here we demonstrate automation of droplet microfluidics based on an inexpensive liquid handling robot for the automated production of human scaffold-free cell spheroids, using pipette actuation and interfacing the pipetting tip with a droplet generating microfluidic chip. In this chip we produce highly mono-disperse 290μm droplets with diameter CV of 1.7%. By encapsulating cells in these droplets, we produce cell spheroids in droplets and recover them to standard formats at a throughput of 85000 spheroids per microfluidic circuit per hour. The viability of the cells in spheroids remains high after recovery only decreased by 4% starting from 96% after 16 hours incubation in nanoliter droplets. Scaffold-free cell spheroids and 3D tissue constructs recapitulate many aspects of functional human tissue more accurately than 2D or single cell cultures, but assembly methods for spheroids, *e.g.* hanging drop micro-plates, has had limited throughput. The increased throughput and decreased cost of our method enables spheroid production at the scale needed for lead discovery drug screening and approaches the cost where these micro tissues could be used as building blocks for organ scale regenerative medicine.

## Introduction

Microfluidics has developed a plethora of applications, particularly in the life sciences and has been part of miniaturizing, increasing throughput and lowering the cost of a large number of macro-scale sample processing and analysis techniques as well as making possible new processes, relying on micro-scale phenomena. Microfluidic technologies have seen some general automation approaches such as pressure-controlled valving^1^, and programmatically reconfigurable devices^2^. However, most of these have required substantial infrastructure outside of the microfluidic device^3,4,5,6,7^ and only very few, such as, *e.g.* the Fluidigm integrated fluidic circuits can be readily interfaced with general laboratory robotics such as liquid handling robots.

A major microfluidic technology platform is droplet microfluidics. It is a high-throughput sample processing and analysis method enabled by miniaturizing the reaction vessels to nano- or pico-liter droplets^8^. Droplet microfluidics relies on structured compartmentalization of immiscible liquids, *e.g.* water and oil in the form of a water-in-oil emulsion stabilized by a surfactant^9,10^. These tiny cell-scale compartments produced with high size accuracy, allow for millions of individual biological events^11^ or chemical reactions to take place simultaneously^12^, mostly chemically and physically separated^13^. Various microfluidic chip types are used in droplet microfluidics to form, merge, inject into, analyze, and sort millions of droplets quickly and efficiently^8^. In biology, droplet microfluidics is used in multiple applications including metabolic pathways interaction studies^14^, single-cell genomics^15,16^ and transcriptomics^17^, digital droplet PCR^18^, direct microbe evolution^19^, as well as cell spheroid formation including cell aggregation in confined environment^20,21^ or cell encapsulation in hydrogels^22^. Immense droplet production rates as high as 1 trillion(!) per hour have been achieved by parallelized droplet generating devices containing up to 10000 droplet generation nozzles.^23,24^

Automation of microfluidics in general and droplet microfluidics, in particular, is an obvious next step for the technology development in this field. It is even more timely now, as more and more commercial applications are being based on droplet microfluidics. Automating even simple steps that are performed on a daily basis in droplet microfluidic workflows - such as droplet generation, as well as sample recovery/droplet breaking - is not a trivial task. For the automation to truly succeed, it has to rely on as simple as possible widely accessible, standardized solutions.

Liquid handling is an essential part of many experiments related to life science; therefore, an abundance of various standardized solutions for liquid handling has already been developed, those include a pipette and robotic pipetting workstations (also known as liquid handling robots). Robots can work without fatigue, increase the throughput, perform consistently, and ensure accuracy and precision^25^.

The pipetting robot has never been used as the only tool to perform all the tasks needed for successful droplet generation and the recovery of their living content. In this paper, we present a fully automated workflow powered solely by a liquid handling robot paired with droplet microfluidic component. As a demonstration of the approach, we use it to automate the production of three-dimensional (3D) cell culture models, cell spheroids.

The use of cell spheroids and other 3D tissue models are emerging areas of interest both in fundamental research as well as in industrial applications, as they provide more predictive *in vitro* models to study fundamental cell biology, disease pathophysiology, and the identification of novel therapeutic agents^26,27^. Spheroids are also considered a central component of regenerative medicine being one of the vital elements of the rapidly growing personalized regenerative medicine field^28,29^.

As the demand for three-dimensional cell culture models increase in the academic, medical and industrial environments, there is a growing need for affordable automation of spheroid production as the process will eventually play a pivotal role in lowering the total expenses of spheroid fabrication. The increased throughput and decreased the cost of our method enables spheroid production at the scale needed for lead discovery drug screening and approaches the price where these micro-tissues could be used as building blocks for organ scale regenerative medicine.

## Materials and methods

### Droplet-generating microfluidic chip

A droplet microfluidics circuit for production of 12.8nl droplets was designed in AutoCAD (ESI Fig. 1). The flow-focusing droplet generating circuits were fabricated from a 2mm thick poly(methyl methacrylate), PMMA plastic sheet using CNC micro-milling (Roland MDX-40A) with a 200μm milling tool. All the micro-channels including the nozzle were 200μm wide and 100μm deep. The inlets and the outlet were drilled with a 1 mm drill by CNC micro-milling machine. Then the chip was cleaned with a detergent using a cleaning brush and carefully washed with water and Milli-Q water to remove any plastic residues from all micro-channels. Subsequently the chip was dried with compressed air. Finally, the chip was sealed with a pressure sensitive adhesive tape (PSA, ARcare® 90445Q), by peeling off one side of the tape and sticking it to the side of the PMMA chip that contained the open micro-channels.

**[Fig. 1:].**
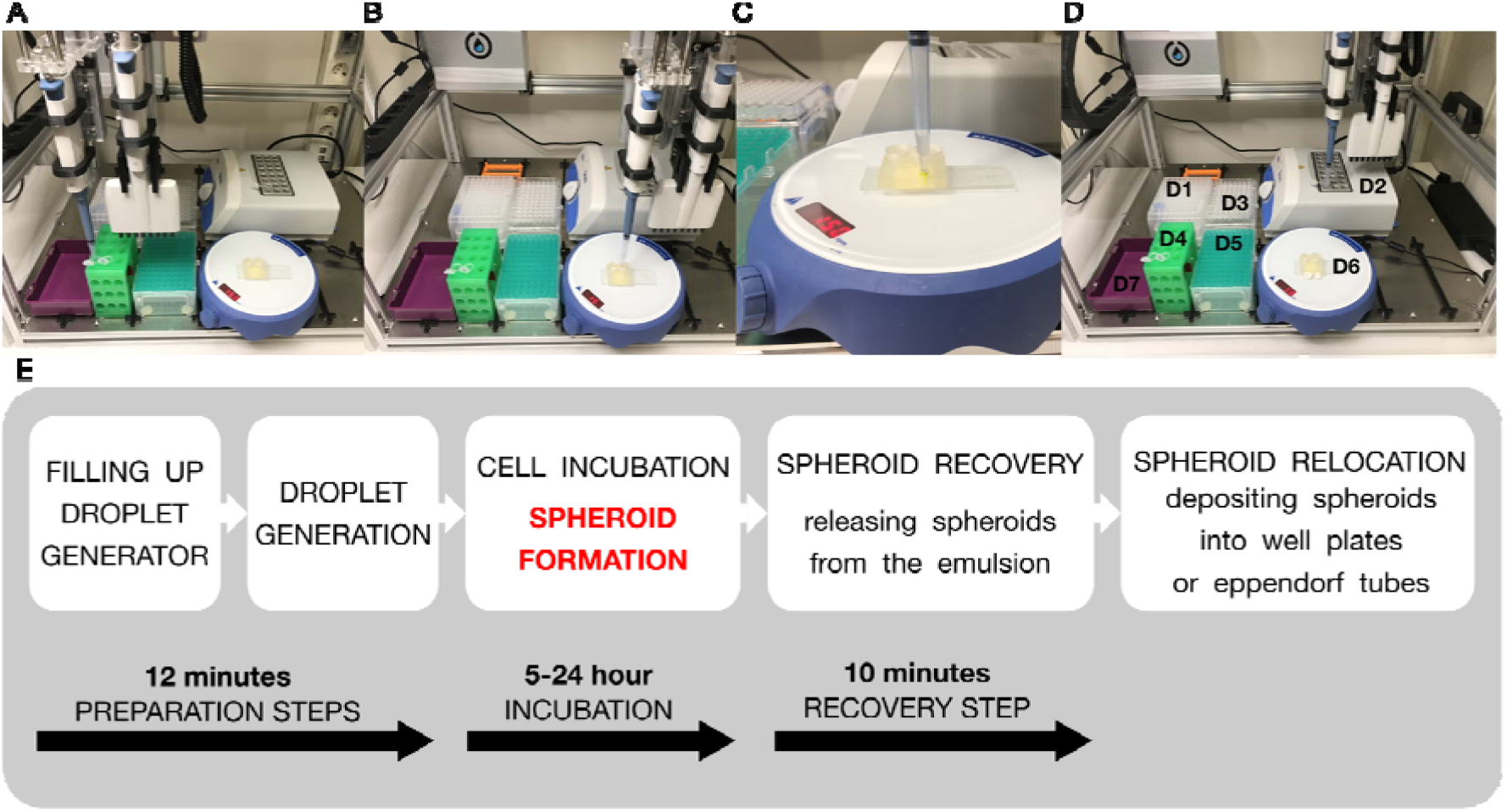

Then a custom-made 3D printed reservoir element was carefully aligned and attached to the other side of the microfluidic droplet generating chip also using an adhesive tape (TESA, top of the droplet generating chip). The tape was pre-cut - 3 holes were cut out using a 2mm Harris Uni-Core biopsy puncher. The holes in the tape were aligned to meet the inlets and outlets of the droplet generating chip and also corresponding outlets and the inlets of the sample and oil reservoir of the 3D printed part.

Finally, the surface of all micro-channels (made from PMMA and PSA tape) was treated with Aquapel. The surface modifying chemical was removed from the channels by compressed nitrogen after 2 min. That step was followed by passing the HFE oil (HFE, 3M™ Novec™ 7500 Engineered Fluid) through all micro-channels, to wash out any remaining of Aquapel. Finally, all the liquids were removed from the chip by compressed nitrogen.

To prevent sedimentation of the cells in the dispersed phase/sample reservoir and clogging the microfluidic micro-channel, as well as for keeping as much as possible equal number of cells entrapped in individual droplets, the whole droplet generating chip was placed on a magnetic stirrer (IKA® big squid) and a sterile small magnet was placed in the dispersed phase/cell sample containing reservoir. The magnetic stirrer was set at 150 rpm.

For collecting droplets, a modified 1ml pipette tip was used (BRAND®, 50-1000μl). A simple 4 millimeter latex sleeve (Kent Elastomer Products, Inc., natural rubber latex tubing with OD 2.3 mm, ID 0.8 mm) was secured at the thinner end of the pipette tip. This rubber sleeve plays an important role, as it seals the connection between the output pipette tip being filled with generated droplets and the outlet of the droplet generating chip.

### Cell sample

HEK293 cells were cultured in FreeStyle293 culture medium in a non-adhesive cell culture flask (Corning® 125ml Erlenmeyer flask, non-pyrogenic, polycarbonate) in a cell culture incubator (Thermo Electron Corporation Hera cell 150, 37 °C, 5% CO2). Before encapsulating in droplets, the cell suspension was prepared to meet the 3.7×10^6^ cells/ml criteria (in FreeStyle293 culture medium).

### Droplet generation process

As a continuous phase a hydrofluoroether (HFE, 3M™ Novec™ 7500 Engineered Fluid) with 1% PEG-PFPE amphiphilic block copolymer surfactant (Ran Biotechnologies) was used. 1ml of HFE with 1% of the surfactant was added to the continuous phase reservoir with the aid of a pipetting robot (Opentrons OT-One-S Hood). This primes the micro-channels, limiting to the minimum the amount of air trapped inside the network of microfluidic channels. The initial step was followed by filling up the dispersed/sample reservoir with 500μl of the cell suspension. Then a negative pressure was applied to the upper end of the droplet collecting element (a modified 1ml pipette tip) by a computer-controlled pipette which the liquid handling robot is equipped with (DragonLAB Top Pipette 100-1000μl). The pipette was programmed to transfer 1000μl in 100 steps (10μl each), at a frequency 0.5Hz. Once applied, the suction force generated by the removes all the remaining air trapped in the system, and starts the negative pressure driven micropipette-based droplet generation process.

### Droplet content recovery

To recover the cells from within the droplets, that is to separate the continuous phase from the dispersed phase a polytetrafluoroethylene, PTFE filter membrane (Sartorius Biolab Products PTFE filter 0.45μm pore size) was used. About 400μl of the emulsion (containing ca 17000 droplets) was placed using a pipetting robot on the surface of the PTFE filter membrane. Immediately, the HFE oil drained into the membrane, destabilizing and breaking the emulsion and allowing the droplets to merge, forming a large droplet full of spheroids floating freely in the culture medium. Alternatively, a chemical-based sample recovery method was used, where 400μl of droplets were merged using 30μl of perfluorooctanol^30,31^. The recovered spheroids were subsequently analyzed for viability.

### Cell viability test

#### HEK cell culture

Before droplet generation, HEK cell culture was analyzed with BIO-RAD TC20TM Automated Cell Counter using BIO-RAD counting slides with a dual-chambers (#145-0011). The cells were stained with Trypan Blue solution (Sigma Life Science, T8154-100ML) which selectively color dead cells in blue. Live cells with intact cell membranes are not colored.

A 10μl of the cell suspension was mixed with 10μl of Trypan Blue and finally 10μl of the mixture was pipetted into one of the chambers of the counting slide. The automatic measurements were triplicated and an average value of viable cells was calculated.

The viability of the used cells was 96% - with a total number of cells reaching 3.88×106/ml and 3.7×106/ml alive, respectively.

#### HEK cell spheroids

To analyze the viability of the cell spheroids that were formed in droplets, a combination of Hoechst 3334 (Life technologiesTM, C47198), Propidium iodine (Life technologiesTM, C27858) and Calcein AM green (Sigma Life Science, 56496-50UG) was used. 1 μl of each was added to 1000μl of 1X PBS. Then, a 100μl of the Hoechst 3334, Propidium iodine and Calcein AM mixture was added to a 50μl solution containing the recovered spheroids (cell aggregates suspended in FreeStyle293 culture medium). This was followed by an incubation step, 30 min in the dark. Finally, fluorescent images of the stained spheroids were taken using fluorescent microscopy with Nikon Eclipse Ti to measure the final viability of formed spheroids (in bright field, FITC, TRITC and DAPI). The number of dead cells was counted in a randomly selected image containing 40 spheroids.

The viability of the cell spheroids (%) was calculated using the formula: number of dead cells/estimated total number of cells forming a spheroid*100.

On average, the viability of the HEK cells forming spheroids was 92,05%.

### Robot’s operations

A. Setup: the robot’s head equipped with 1ml pipettor grabs a regular 1ml pipette tip from the tip rack (Fig. 1D_1) by locking the pipettor’s lower end into the pipette tip - just like a human would do - and moves it towards the Eppendorf tube that contains HFE oil supplemented with the surfactant (Fig. 1D_4). Then it lowers itself into the vessel and aspirates 950μl of the HFE oil and transfers it into the continuous phase (smaller) reservoir of the droplet generating microfluidic chip (Fig. 1D_6). The robot’s head then moves back to the initial position (Fig. 1D_1) and returns the tip into the same well of the tip rack, so that it could be used later for the same purpose. Similarly, but with a new pipette tip collected from a neighboring position, the robot transfers 450μl of the cell sample to a slightly bigger reservoir (dedicated to the dispersed phase), that contains at this point also a rotating magnet. Once the sample is pipetted out, the robot moves its head towards the waste box (Fig. 1D_7), and drops contaminated tip there.

B. Droplet generation: from the A1 position of the 1ml pipette tip rack (Fig. 1D_1), the robot collects a modified pipette tip that has a seal (an O-ring) and moves its head above the outlet of the droplet generating microfluidic chip. Before reaching its destination, it drops the plunger of the pipettor to the minimum removing any remaining air from the pipette’s pumping mechanism. Then it lowers itself, reaching the level of the actual outlet of the droplet generating microfluidic chip (the PMMA part, Fig. 1C, and 2C). The connection between the pipettor and the microfluidic chip becomes truly sealed when the bottom of the O-ring reaches the level of PMMA and compresses against it. At the same time, the conical shapes of both, the pipette’s tip covered with the rubber seal and the 3D printed well locks, enhancing the connection by making it air- and liquid-tight. Then the controlling software activates generation of the negative pressure in the microfluidic system by sequentially lifting the plunger and aspirating 10μl with a frequency of 0.5 Hz, every 2 seconds. The robot repeats this step 100 times. It is essential to generate negative pressure gradually, for example, by a sequential lift of the plunger, as this helps in maintaining a sealed connection between the rubber O-ring and the microfluidic chip. Otherwise, the air-tight connection might fail as the vigorously sucked air bursts through the seal preventing a constant build-up of the pressure difference between the pipettor and the 3D printed reservoirs. If that happens, bubbling inside the pipette tip is observed and droplets are no longer generated since not strong enough negative pressure is transferred through the channels of the microfluidic chip^33^. Both of the liquids, the cell suspension, and the HFE oil are passed through a network of separate microchannels until they reach the flow focusing part of the microfluidic chip. There, the cell suspension (dispersed phase) is being “pinched away” by the HFE oil (continuous phase) forming the droplets with a similar number of cells (Fig. 3A, B)^34,40^. A fluorosurfactant was used to stabilize the emulsion at a concentration of 1% (w/v) and was dissolved in the continuous phase before the experiment.

**[Fig. 2:].**
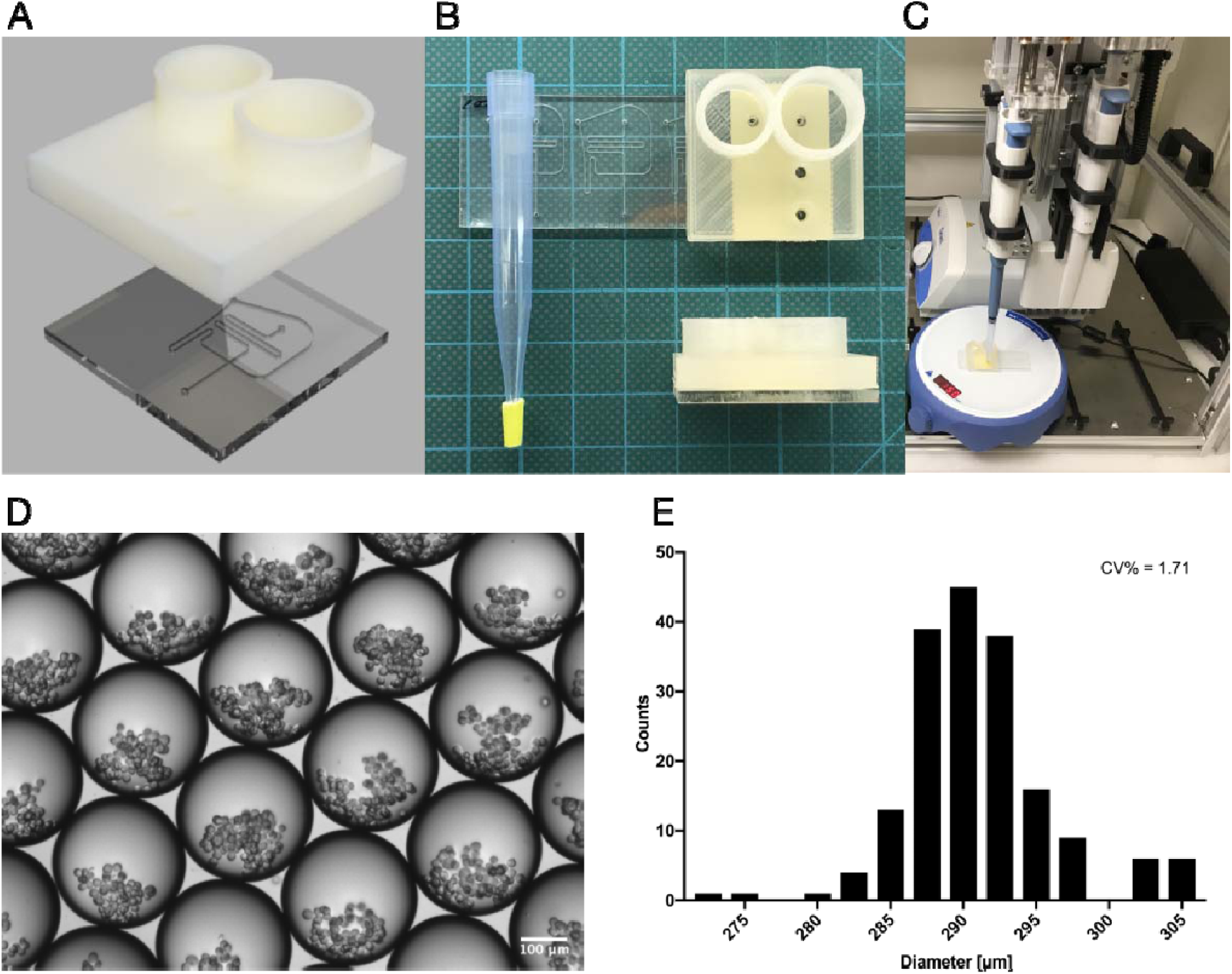

After this step, the robot pauses for 300 seconds until a full one milliliter of liquid transfers through the network of the microchannels into a modified pipette tip. The pause in the workflow is needed for the pressure difference to equalize between the pipettor and the atmospheric pressure through the network of microchannels of the chip^33^. The pause limits also sucking-in an excess amount of air into the droplet-containing pipette tip, which prevents the formation of a polydisperse emulsion.

Now, the tip attached to the pipettor doubles as a temporary droplet reservoir. Once all the droplets are collected, the robot lifts its pipettor and moves it above the heat block. The head is lowered, and the droplets are transferred into an Eppendorf tube where the cells will be incubated for at least 16h and will transform into spheroids. To prevent evaporation of droplets during the extended incubation, the emulsion is stored in Eppendorf tubes under a protective, immiscible and non-volatile layer formed by mineral oil.

C. Spheroid recovery: once the overnight/16h lasting pause is finished the robot activates itself and grabs a clean tip from the 1ml pipette tip rack (Fig. 1D_1). Then, depending on the method chosen for the droplet recovery it travels to the Eppendorf tube rack (Fig. 1D_4) and transfers 30μl of the droplet breaking chemical (perfluorooctanol) into an Eppendorf tube, or it moves directly to the heat block. At the heat block site, the robotic arm lowers 1ml pipettor into an Eppendorf tube in which the droplets were incubated overnight and collects its content (1000μl), leaving - if present - the mineral oil behind. Then the robot pauses for an additional 60 seconds, allowing the droplets to cream at the top of the HFE oil (now stored again in the pipette tip).

Finally, the robot lifts its head a few millimetres up and removes 600μl of the HFE oil that remains in the lower part of the tip, saving only the packed emulsion. Then, depending on the selected recovery method, the robot moves the pipettor either to the Eppendorf tube rack (Fig. 1D_4) or to a neighbouring position which has a recovery membrane.

## Results and discussion

The robotically-automated droplet microfluidic platform consists of four major components: a liquid-handling robot, a droplet generating microfluidic chip, an interface connecting them and a computer that controls every movement and action. The platform is built on top of a commercially available liquid-handling robot. Opentrons OT-One Hood was chosen because of its open source, simple design and relatively low price ($4000). The Opentrons OT-One Hood robot is essentially a standard 1ml or 8 channel 300μl pipettor attached to the robotic arm, that can move in all directions according to simple Python code. The liquid handling robot can be controlled either by a Python script or “manually” using a Python console and XYZ coordinates, as well as using Opentrons own software - in all the cases an external computer with appropriate open source software has to be connected to the robot.

Although the robot has 15 slots for the well plates on its deck, the design of the device allows accommodating only 8 of them, meaning that only part of the surface of the deck (about 30×40 cm) can be used for fully automated operations. The rest of the surface is out of reach for both or just one of the pipettors attached to the robotic arm.

To fit the necessary equipment, a 1ml tip rack, a waste box, a 12 slot Eppendorf tube rack, a 96-well plate, a full-size magnetic stirrer with a droplet generating microfluidic chip on top of it, and a heat block for the Eppendorf tubes, we have used all of the 8 available slots.

To generate relatively large quantities of highly mono-disperse nanoliter droplets a source of pressure (negative^32,33^ or positive^34,35^) and a microfluidic chip based on specifically organized microchannels is needed^36^. To fulfill those requirements, we chose a simple flow-focusing design as the droplet generating element, which we fabricated in a 2 mm thick poly(methyl methacrylate), PMMA plastic sheet using a CNC micro-mill^37^. The CNC engraved PMMA sheet was sealed with pressure sensitive adhesive tape, forming a closed network of microfluidic channels. All microchannels including the nozzle were 200μm wide and 100μm deep (Fig. 2A). To minimize the complexity of the platform, we have decided to actuate the microfluidic device using the 1ml pipette which the robot is equipped with^33^. In this way we eliminate the need for expensive syringe pumps or pressure controllers as the means of actuating liquids in droplet generation. This decision also greatly simplified the control protocol, as only a single device, the liquid handling robot equipped with 1ml pipette needs to be programmed. To be able to use a pipettor as a negative pressure source for droplet microfluidics^33^ we had to devise an interface that connects the pressure source, the pipettor and pipette tip, with the microfluidic chip without losses in suction force generated by the pipettor. Thus, we had to seal the connection between the lower opening of the pipette tip and the outlet of the microfluidic channel.

In the case of microfluidic chips fabricated from polydimethylsiloxane (PDMS), the polymer itself can act as a seal^38^. Unfortunately, PDMS chips lack of reproducibility as those elements are frequently manufactured in low quantities, usually directly by hand - introducing a great human error. And reproducibility is the critical element of successful lab work and the automation^39,4^.

In order to connect seamlessly 1ml pipette tip with the outlet of the microfluidic channel we have added an approximately 4 mm long rubber O-ring at the lower opening of the pipette tip. The conical shape of the 1ml pipetting tip prevents the rubber O- ring from sliding along the plastic element and locks it in place. This modification allows the robot to use the same modified pipetting tip multiple times in a series of workflows (Fig. 2B). To further increase the quality of the connection between the negative pressure source - the pipettor and the outlet of the microfluidic chip, we have added a unique 3D printed part with a 5 mm deep and 3 mm in diameter well (and two reservoirs) on top of the plastic microfluidic chip.

The 3D printed part consists of two other crucial elements - two reservoirs for both the continuous and the dispersed phase - the HFE oil supplemented with 1% of a surfactant and the cell sample, respectively (Fig. 2A). We have dedicated a smaller reservoir to the continuous phase, as its volume is 1.15ml, which support successful droplet generation using 1ml pipette. A slightly greater reservoir, with a capacity of 1.65ml, has dimensions that allow the presence of a small magnetic stirrer submerged in the cell sample to prevent the sedimentation of the cells during the process of droplet formation (Fig. 2B).

For the robot to operate smoothly, the device has to go through the initiation round. The operator has to make sure that the 1ml pipette tip rack (Fig. 1D_1) has clean tips loaded. The first row of the rack should have a modified pipette tips (equipped with a special rubber O-ring). The heat block (Fig. 1D_2) has to be switched on and set at 37°C. Depending on the number of samples, one or several Eppendorf tubes filled with 0.5ml of mineral oil have to be placed in the heat block’s well to receive droplets at certain point of the protocol. A 96-well plate (Fig. 1D_3) or an additional Eppendorf tube has to be placed on the deck or in an Eppendorf tube rack (Fig. 1D_4), respectively - here the formed and recovered spheroids will finally be deposited. Then an Eppendorf tube with the HFE oil supplemented with 1% of surfactant should be located in the first (A1) position of the Eppendorf tube rack (Fig. 1D_4), followed by another one with the cell sample in the neighboring (B1) position of the same tray (Fig. 1D_4). Finally, the magnetic stirrer (Fig. 1D_6) should be centered under the microfluidic chip. Lastly, a small PTFE coated magnet should be placed into the cell sample reservoir and the magnetic stirrer should be switched on.

The automated workflow for generating cell spheroids using robotized droplet microfluidic platform consists of five steps: sample and reagent loading, droplet generation, spheroid formation, cell incubation, spheroid recovery, and spheroid dispensing. All steps are performed independently by the liquid handling robot, without assistance from an operator.

The liquid handling robot throughout all of those steps follows exactly the experimental protocol uploaded in the form of a Python script to the controlling computer. It automatically transfers samples, generates, collects and deposits droplets using the standard and modified pipette tips, respectively.

Even though a regular 1ml pipette operated by a robot was used as a negative pressure source, the measured size distribution of generated droplets (CV%) was 1.71. The generated droplets with the cells inside them were very similar in size (Fig. 2E).

### Cell incubation/spheroid formation part

It is important to note that while Opentrons robots are not able at this point to integrate easily with a proper external incubator designed specifically for cell culture, other - more expensive - liquid handling robots can, allowing for the spheroid formation part of the protocol to be performed in an external incubator. Also, if there is a need, simply the Eppendorf tube with the emulsion can be manually relocated to an external incubator for the overnight incubation. In that case, the protective mineral oil layer should not be used as the cell culture incubators are humidified. We have tested both paths, where the cells in droplets were incubated in an external incubator but also on the heat block in an Eppendorf tube, where the surrounding droplets HFE oil was used as the gas reservoir for the spheroid forming cells^20,21,40^, and both methods works.

While the robot is inactive, the cells are stored at 37°C. As noted elsewhere it takes several hours for the cells to form loose aggregates that finally convert into spheroids^20,21,40^. The spheroids are essentially tight cell aggregates that do not fall apart when exposed to an external force^41^. In our case, a properly formed spheroid should survive the process of sample recovery as well as physical relocation performed by a pipetting robot. Because our method does not rely on any hydrogel that may maintain the cell aggregates together^40,42,43,44^ without a proper biological integration and a physical interaction of the cells, the aggregates fall apart the moment the droplets are broken, and the physical micro-compartmentalization disappears. Since our platform does not allow for the real-time observation of the biological processes that appear in droplets during the spheroid formation part, to explain what happens in the droplets, we have to rely on the hypothesis that was given by Baroud et al.^40^ on how the cells sediment and aggregate into spheroids in droplets. It is worth noting that although our platform uses similar droplet size it does not lock the droplets in wells, and by that, it does not prevent the HFE oil to freely flow around the whole droplets, allowing the flow to be transferred through the water-oil interface into the droplets. This additional internal flow could potentially change the behavior of the cells in droplets and at least delay their sedimentation.

The minimum time needed for the cells to aggregate efficiently enough, so they maintain the spheroidal form after the recovery was about 5h, which is in line with the results that other groups were observing^20,40,45^.

Due to convenience, we have decided that the cells will be incubated overnight (16h), giving the cells enough time to interact with each other so they could form a stable spheroid.

#### A chemical-based sample recovery

In the case of the chemical-based sample recovery method, 400μl of the packed emulsion is pipetted into an Eppendorf tube that already contains 30 μl of perfluorooctanol, which has been pipetted there right after the 16h pause was finished (Fig. 1D_4). Then the robot automatically changes the pipetting tip for a new one and initiates a 5-minute pause required for the perfluorooctanol to break the emulsion entirely and for the droplets to merge into one that contains all of the spheroids. Once the break is finished, the robot aspirates 500μl from the spheroid comprising Eppendorf tube (to be sure that the whole of the sample is collected). This step is followed by pipetting out 125μl, which contains the air and all of the waste (the HFE oil mixed with perfluorooctanol). At this point the pipettor’s tip holds all of the recovered spheroids suspended in the culture.

#### A membrane-base sample recovery

If the membrane sample recovery method is chosen, then the robot transfers the remaining 400μl of the packed emulsion onto a PTFE membrane that is located next to Eppendorf tube rack (Fig. 1D_8). The droplets spread around the membrane and eventually the two phases separate from each other.

Then the robot automatically changes the pipetting tip for a new one and initiates a 5-minute pause required for all of the droplets to merge. After 5 min., the robotized pipettor collects all spheroids by aspirating directly from PTFE membrane.

Finally, the robot enters into the sample relocation step. Here, it optionally dilutes the spheroid containing sample with culture medium and if programmed, distributes 75μl segments of the sample into a preselected - in the Python script - wells of the 96- well plate (Fig. 1D_3). This step finishes the automated workflow, which can be now repeated with a new set of samples.

#### The mechanism of the novel sample recovery method based on a physical phase separation on a PTFE membrane

The process of membrane-based sample recovery described above is constructed on the physical separation of two phases by oil-attracting (oleophilic) and a water-repelling (hydrophobic) membrane made of polytetrafluoroethylene, PTFE^46^. Similar techniques based on a membrane separation of immiscible liquids are already known in the field of oil and water separation^46,47,48^. This process is also described as de-emulsification and is essential in the industries involving the recovery of solvents and desalination of the oils as well as in the petroleum industry, especially during crude oil production^49,50,51^.

It has been shown before that the membranes made of PTFE or composites membranes based on PTFE as well as other hydrophobic-oleophilic membranes are incredibly useful in filtering out the oil from water^46,47,48,52^. To our knowledge, this phenomenon has never been applied in the field of droplet microfluidics especially for a chemical-free biological sample recovery from the emulsion.

The success of the method is centered around a substantial difference in the contact angle with the PTFE membrane of the two components that form the emulsion. The contact angle of the HFE-7500 oil used in our experiments as the continuous phase with PTFE has been reported to be 0° ^53^. On the other hand, the water contact angle on the same type of membrane ranges between 90 and 110° depending on the pH^54^. A contact angle close to zero means that the surface has oleophilic properties. Whereas on a hydrophobic material water forms a droplet with a contact angle greater than 90° ^55^.

Once the emulsion is deposited on top of the PTFE membrane, the oleophilic nature of polytetrafluoroethylene attracts its oil component. The HFE oil wets the surface of the PTFE and spreads around and across the membrane immediately after the emulsion hits its surface (Fig. 4A). With the physical removal of the HFE oil also comes the withdrawal of the surfactant. Those two factors destabilize the emulsion forcing the droplets to merge (Fig. 4B, 4C). This process is rapid and straightforward.

**[Fig. 4:].**
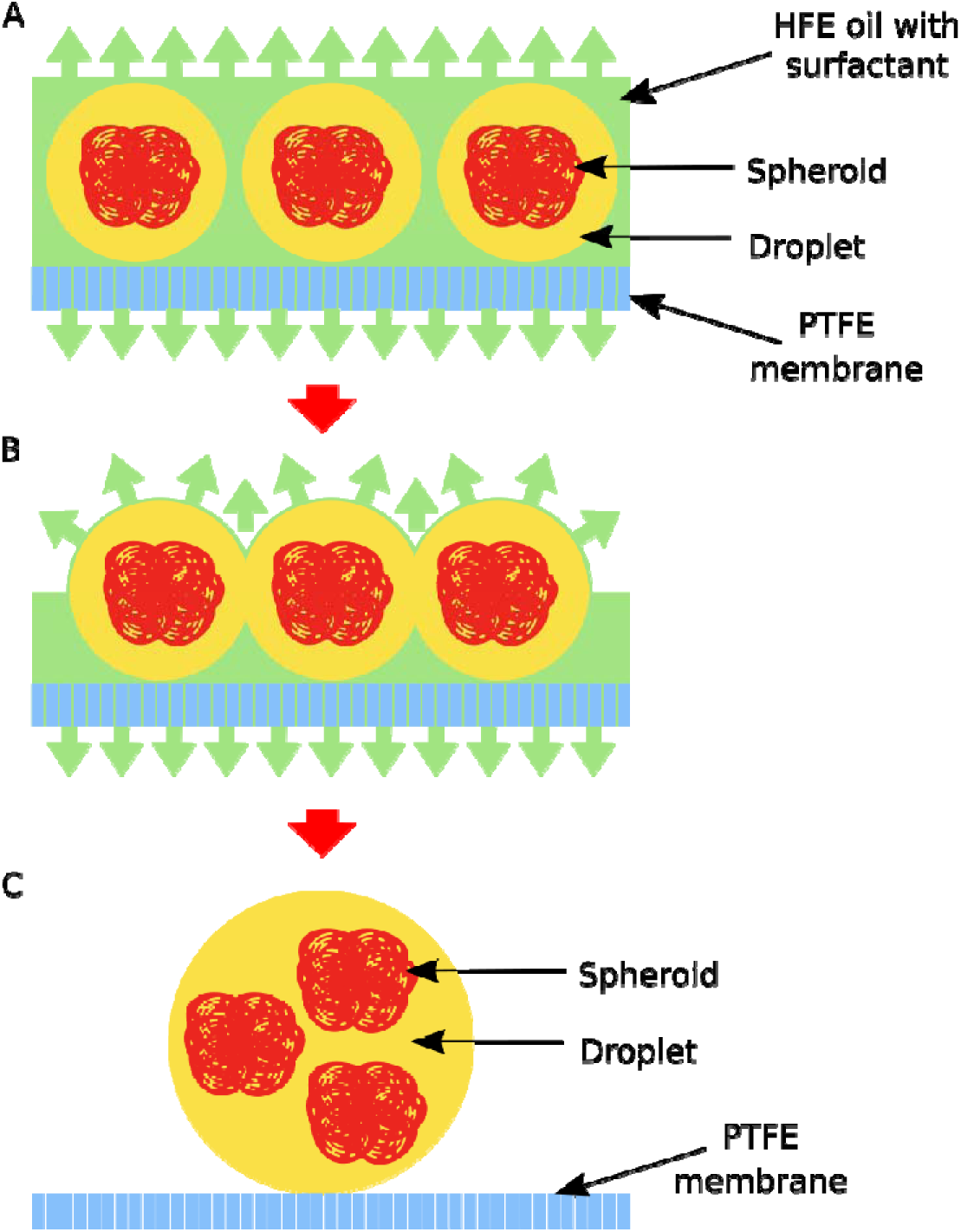

The new method presented here allows for a simple chemical-free sample de-emulsification for droplet microfluidics, which is inert against the cells^56^. Others have already proven that commonly used chemicals such as perfluorooctanol may interfere with the results of the biological experiments performed in droplets^57^.

It is worth noting that the new method described above has all of the three components: is easy to scale up and is ready for automation, at the same time being neutral towards the biological content of the sample entrapped in droplets.

### Spheroid viability

The results from other groups indicate that droplet microfluidics used for spheroid formation has limited impact on the viability of the cells entrapped and the spheroids formed in droplets^20,40,45^. The majority of already published droplet microfluidic-based spheroid forming protocols rely on double emulsion setups^20^ or on adding hydrogels^40,42,43,44^ to the culture medium in which the cells aggregate.

It was unknown to us how the phase separation technique (membrane-based sample recovery) used for releasing the dispersed phase from the emulsion influences the overall viability of the cells that form spheroids.

To test the cells viability a 100μl of the Hoechst 3334, Propidium iodine and Calcein AM mixture was added to a 50μl of the recovered spheroids (cell aggregates suspended in FreeStyle293 culture medium that they were formed in). This step was followed by 30 min incubation in the dark. Finally, fluorescent images of the stained spheroids were taken (in FITC, TRITC and DAPI, Fig 3C) and the number of dead cells was counted on a randomly picked image with 40 spheroids on it.

**[Fig. 3:].**
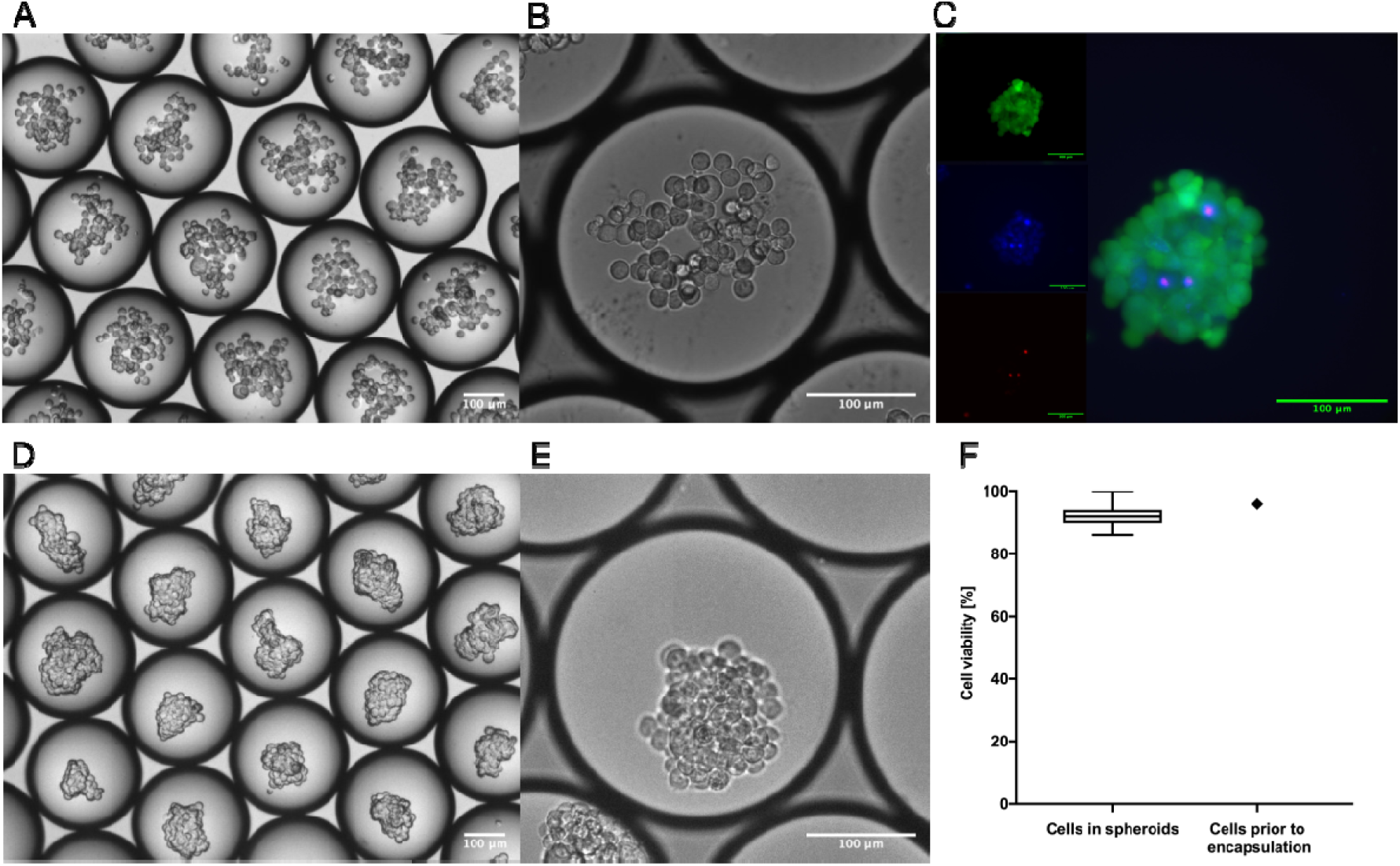

The overall viability of the sample after an overnight incubation during which the cell aggregation took place and the cell spheroids were formed was 92% (Fig 3F). The initial viability of the cells used to form spheroids was 96%, indicating that the whole process of the transformation of cell suspension into cell spheroids using our automated droplet microfluidic platform resulted in a 4% drop in cell viability, which is in line with the results that other group presented when droplets were used to form spheroids^20,40,45^.

### HEK cell spheroids produced using automated droplet microfluidics

We have analyzed the circularity of the spheroids using ImageJ’s measure command that computes circularity of an object according to a formula: 4π*area/perimeter^2^, where circularity of 1.0 indicates a perfect circle^46^. The mean value of circularity of analyzed spheroids was 0.699 with a standard deviation of 0.0695. Also, the mean value of the aspect ratio of spheroids was 1.369 (Std. dev. 0.252). Both parameters indicate that the spheroids are not perfect spheres - this complies with the images taken (Fig. 3D). This variability could be due to cell aggregation before droplet formation which was also reported already^40^. According to theory, droplets form the closest packing of undeformed spheres at a volume fraction of 0.7405^47^. Based on that estimation, about 26% of the densely packed emulsion generated by the automated droplet microfluidic platform was the continuous phase (HFE oil), and the remaining 74% was the dispersed phase - the spheroids suspended in culture medium. Knowing that a single droplet has a volume of 12.76nl and that all of the generated droplets contain a similar number of cells, we can estimate that in one run robotically-automated droplet microfluidic platform can produce over 17000 of droplets/spheroids. Since the duration of one-run is 12 minutes, in 1 hour the robot can cycle five different samples generating up to 85000 spheroids. In theory, at this stage of the development, the maximum capacity of the platform is 120 different samples, which would result in producing 2.04×10^6^ spheroids in 48 hours, as similar time is needed for the robot to recover the spheroids from the emulsion after overnight incubation.

## Conclusions

A robotically-automated droplet microfluidic platform presented in this paper allows for independent from the human operator, therefore automated, production of cell spheroids. The new platform was tested as a self-sufficient assembly line for scaffold-free HEK cell spheroids, allowing generation of over 17000 spheroids in one run. The system’s maximum throughput was estimated at 85000 spheroids an hour.

A new scalable method for sample recovery from the droplets, which eliminates the need for the use of chemicals for disrupting the emulsion supports the automation process.

We believe that the low price of the main component of the robotically-automated droplet microfluidic platform and high production throughput, and full automation of the process decreases the generation cost of a single spheroid to the scale needed for it to lead drug discovery screening, matching the costs where spheroids could be used as building blocks for an organ scale regenerative medicine.

